# Loss of Farnesoid X Receptor Signaling Promotes Choroidal Neovascularization in Mice

**DOI:** 10.64898/2026.10.21.760483

**Authors:** Chanel Harris, Mohammed A. Nisar, Ankit Seth, Allston Oxenrider, Briah Bailey, Gayatri Seth, Pamela M. Martin, Ravirajsinh N. Jadeja, Menaka C. Thounaojam

**Affiliations:** Department of Biomedical Sciences, School of Graduate Studies, Meharry Medical College, Nashville, TN, USA; Departments of Cellular Biology and Anatomy, Medical College of Georgia at Augusta University, Augusta, GA, USA

**Keywords:** Farnesoid X receptor, choroidal neovascularization, age-related macular degeneration, angiogenesis, fibrosis

## Abstract

Choroidal neovascularization (CNV) is a defining feature of neovascular age-related macular degeneration (nAMD). Although anti-vascular endothelial growth factor (VEGF) therapies suppress neovascular activity, incomplete responses, recurrent leakage, and subretinal fibrosis remain important clinical challenges. The farnesoid X receptor (FXR) is a bile acid–activated nuclear receptor with anti-inflammatory, anti-fibrotic, and anti-angiogenic functions in other tissues, but its role in CNV is unknown. Analysis of publicly available human transcriptomic data (GSE135092) showed FXR expression in the retinal pigment epithelium (RPE)/choroid and significant downregulation in AMD-affected macular RPE/choroid. To define the role of FXR in CNV, laser photocoagulation was performed in wild-type (WT) and FXR-deficient (FXR^−/−^) mice, with disease progression assessed by in vivo imaging, histology, *ex vivo* choroidal sprouting, and endothelial cell assays. FXR expression in the RPE/choroid was reduced 14 days after laser injury. Compared with WT mice, FXR^−/−^ mice developed larger CNV lesions, more extensive hyperfluorescence on fluorescein angiography, and increased lesion area during disease progression. FXR deficiency also impaired resolution of fibrotic remodeling. In WT mice, α-SMA peaked at day 14 and declined by day 21, whereas collagen I remained low throughout; in FXR^−/−^ mice, both collagen I and α-SMA were significantly elevated relative to WT and remained significantly higher than WT at day 21. Consistent with these findings, FXR- deficient choroidal explants exhibited increased vascular sprouting, and FXR^−/−^ endothelial cells formed more extensive capillary-like networks. Together, these findings identify FXR as an endogenous regulator of pathological angiogenesis, vascular leakage, mesenchymal activation, and fibrotic remodeling during CNV and provide a rationale for investigating FXR-directed strategies for nAMD.

**Impact Statement:** Neovascular age-related macular degeneration remains limited by recurrent vascular leakage and subretinal fibrosis despite effective VEGF suppression. This study identifies the bile acid-activated nuclear receptor FXR as a previously unrecognized regulator of the angiogenic and fibrotic responses underlying choroidal neovascularization. Integrating human AMD transcriptomic data with genetic, in vivo, ex vivo, and endothelial-cell approaches, we show that FXR expression is reduced in diseased RPE/choroid and that FXR deficiency enhances neovascular growth, vascular leakage, endothelial angiogenic activity, and persistent myofibroblast activation. These findings connect bile acid receptor signaling with RPE/choroidal vascular homeostasis and provide a biological foundation for evaluating FXR-directed approaches as complementary strategies for neovascular AMD.

## Introduction

Age-related macular degeneration (AMD) is a leading cause of irreversible central vision loss among older adults and represents a growing global public health burden ^1^. Among the clinical forms of AMD, neovascular AMD (nAMD) accounts for the majority of severe vision loss despite comprising a smaller proportion of total AMD cases ^2^. The defining pathological feature of nAMD is choroidal neovascularization (CNV), a process characterized by the growth of abnormal choroidal blood vessels through Bruch’s membrane into the subretinal and sub-retinal pigment epithelium (RPE) space ^3^. These immature neovessels are highly permeable, resulting in vascular leakage, hemorrhage, exudation, and progressive disruption of retinal architecture that ultimately leads to vision impairment ^4^.

Although anti-vascular endothelial growth factor (VEGF) therapies have improved the management of nAMD, important limitations remain ^5^. Anti-VEGF agents effectively suppress vascular leakage and neovascular growth but do not completely eliminate CNV lesions ^6^ and fail to adequately prevent fibrotic progression ^7^. Long-term studies have reported that a substantial proportion of patients develop subretinal fibrosis despite continuous anti-VEGF treatment, contributing to irreversible visual decline ^8^. Furthermore, the requirement for frequent intravitreal injections imposes a significant treatment burden, and a subset of patients exhibit incomplete or diminishing therapeutic responses over time ^9^. These observations suggest that additional molecular pathways beyond VEGF contribute to CNV pathogenesis and lesion progression.

The development of CNV is a complex process involving coordinated interactions among retinal pigment epithelial cells, choroidal endothelial cells, inflammatory cells, and extracellular matrix components ^10^. After injury and chronic inflammatory stimulation, activated endothelial cells invade the subretinal space and recruit inflammatory and fibrotic cells. Consequently, fibrovascular scar formation develops beneath the retina, contributing to structural damage and permanent vision loss ^11, 12^. These angiogenic, inflammatory, and metabolic processes, and their modulation by aging and sex, have been extensively characterized in murine models of neovascular AMD ^13^. Therefore, identifying endogenous mechanisms that simultaneously regulate angiogenesis, inflammation, and fibrosis may provide new therapeutic opportunities for nAMD.

Farnesoid X receptor (FXR; NR1H4) is a ligand-activated nuclear receptor ^14^ originally identified as a key regulator of bile acid metabolism in the liver, intestine, and gallbladder ^15^. Emerging evidence, however, indicates that FXR exerts broader biological functions extending beyond metabolic regulation ^16, 17^. FXR signaling suppresses oxidative stress, inflammatory responses, fibrosis, and abnormal vascular remodeling in multiple tissues ^18, 19^. Activation of FXR in non-ocular tissues regulates the expression of genes involved in barrier integrity, angiogenesis, cellular stress responses, and inflammatory signaling pathways, all of which are implicated in AMD pathogenesis ^20^. Moreover, previous work from our group demonstrated that ursodeoxycholic acid (UDCA), a bile acid with FXR-modulatory activity, effectively halts pathological retinal neovascularization in the oxygen-induced retinopathy model, highlighting the therapeutic potential of bile acid signaling pathways in the regulation of ocular angiogenesis^21^. Consistent with a broader role for sterol metabolism in ocular neovascular disease, we have also shown that activation of the cholesterol-metabolizing enzyme CYP46A1 reduces pathological retinal neovascularization in the oxygen-induced retinopathy model ^22^.

Secondary conjugated bile acids like ursodeoxycholic acid and tauroursodeoxycholic acid help protect the RPE ^23^, lower the risk of laser-induced CNV, and reduce VEGF levels ^24, 25^.

Further, patients with neovascular AMD often show changes in their circulating bile acid profiles^26^. While the biological relevance of FXR signaling to the outer retina is supported by these prior studies, the contribution of FXR signaling to CNV development and lesion remodeling has not been investigated.

Based on these findings, we hypothesized that FXR acts as an endogenous suppressor of CNV progression by regulating angiogenic, inflammatory, and fibrotic signaling pathways within the retinal and choroidal microenvironment. Using a laser-induced CNV model, we investigated the effects of FXR deficiency on lesion development, vascular leakage, retinal remodeling, fibrosis, and choroidal endothelial cell angiogenic behavior. Our findings identify FXR as a critical regulator of CNV progression and suggest that therapeutic activation of FXR may represent a novel strategy to simultaneously target angiogenesis, inflammation, and fibrosis in neovascular AMD.

## Materials and Methods

### Analysis of human retina and RPE/choroid transcriptomic data (GSE135092)

The human dataset GSE135092 was obtained from the NCBI Gene Expression Omnibus (GEO)^27^. This dataset includes bulk RNA-sequencing data from 537 tissue samples collected from human donors. The samples represent both the RPE/choroid and the neurosensory retina, spanning macular and non-macular regions, and include both AMD-affected and control donors. Sample-level metadata were extracted from the GEO series matrix file, and raw read counts and normalized expression values were compiled into a unified gene-by-sample matrix for downstream analysis. Because the raw sequencing reads are not available for this dataset, all analyses were performed using the processed read counts and normalized expression values provided in the GEO supplementary files.

Tissue-compartment specificity of NR1H4 was assessed by comparing expression levels between the RPE/choroid and neural retina, using canonical RPE identity markers (RPE65, BEST1, LRAT, RLBP1, TYR) as reference. Differential expression between AMD and control macular RPE/choroid samples was determined using a negative binomial generalized linear model (DESeq2) on raw count data, including all available samples (26 AMD, 105 control; design: ∼ AMD status). P values were adjusted for multiple testing using the Benjamini-Hochberg method.

### Animals

Heterozygous farnesoid X receptor-null (FXR^+/−^) mice (Jackson Laboratory, Stock No. 004144), maintained on a C57BL/6J background, were intercrossed to generate homozygous FXR- deficient (FXR^−/−^) mice and wild-type (WT) littermate controls. Genotypes were confirmed by PCR analysis of tail DNA according to the supplier’s protocol. Male mice were used at 8 weeks of age, with age-matched WT littermates as controls throughout the study. In vivo imaging (fundus photography, fluorescein angiography [FA], and optical coherence tomography [OCT]) was performed longitudinally in the same animals at days 7, 14, and 21. Separate cohorts (n = 5 per genotype per time point) were euthanized at each time point for histological and flat-mount analyses. All animals were housed in a standard experimental facility with free access to food and water. The facility was maintained under a 12-hour light-dark cycle. All procedures conducted throughout the study were in accordance with the Association for Research in Vision and Ophthalmology’s (ARVO) guidelines for the Use of Animals in Ophthalmic and Vision Research and were approved by the Institutional Animal Care and Use Committee (IACUC) of Augusta University (#2022-1083) and Meharry Medical College (#24-06-1486). Mice were euthanized by CO inhalation followed by cervical dislocation, consistent with AVMA guidelines and the approved IACUC protocol.

### Laser-Induced CNV Model

Mice were anesthetized by intraperitoneal injection of ketamine (100 mg/kg) and xylazine (10 mg/kg). Pupils were dilated with 1% tropicamide ophthalmic solution, and a lubricating ophthalmic gel was applied to maintain corneal clarity during the procedure. Experimental CNV was induced using a Micron IV image-guided laser photocoagulation system (Phoenix Research Laboratories, Pleasanton, CA, USA). Both eyes of each animal were treated, with four laser burns delivered per eye at locations surrounding the optic nerve head using a 532-nm green laser with a spot size of 50 µm, a duration of 70 ms, and a power setting of 240 mW. Successful rupture of Bruch’s membrane was verified by the appearance of a cavitation bubble immediately following laser application. After laser treatment, ophthalmic antibiotic ointment was applied to the eyes, and animals were allowed to recover under standard housing conditions ^28^. For OCT, lesions were imaged in vivo in both eyes (4–6 analyzable lesions per animal); for flat-mount immunostaining, 2–4 lesions per animal were analyzed. Lesion measurements were averaged within each animal, and the animal was used as the statistical unit.

### Western blot analysis

RPE/choroid lysates were prepared from untreated control and CNV mice using RIPA buffer (Thermo Fisher, Waltham, MA, USA) supplemented with 1% phosphatase and protease inhibitors (Sigma-Aldrich, St. Louis, MO, USA). Protein (30–50 µg) was resolved by SDS-PAGE and transferred to PVDF membranes. Membranes were blocked with 5% non-fat dry milk and probed with primary antibodies: FXR/NR1H4 (A24015, 1:1000; ABclonal, Woburn, MA, USA) and TGF-β1 (81746-2-RR, 1:1000; Proteintech, Rosemont, IL, USA). Additionally, β- actin (sc-47778, 1:1000; Santa Cruz Biotechnology, Dallas, TX, USA) was used as a loading control. Secondary detection was performed using chemiluminescence (Thermo Fisher) and the Azure 300 Imaging System. Band intensities were quantified using NIH ImageJ software and expressed as fold change relative to controls.

### Choroidal Sprouting Assay

The choroidal sprouting assay was performed as previously described with minor modifications^29^. Briefly, eyes were collected from 21-day-old WT and FXR^−/−^ mice, followed by careful removal of the optic nerve, lens, and cornea. Three-week-old mice were used because choroidal explants from young mice sprout more reproducibly in this assay; the concordant phenotype in adult animals in vivo supports its relevance to the CNV model. The remaining RPE/choroid/sclera complex was isolated and cut into approximately 1.0 mm × 1.0 mm explants. The bottom of each 24-well plate was coated with growth factor-reduced Matrigel (Corning, Corning, NY, USA). The explants were embedded into the Matrigel and placed in the incubator for 10 minutes to solidify. After solidification of the Matrigel, endothelial growth medium-2 (EGM-2; Lonza, Walkersville, MD, USA) was added to each well, and the plates were placed in a humidified incubator with 5% CO at 37°C.

Choroidal explants were monitored for endothelial sprouting over 7 days, and culture medium was replenished every 2 days. At days 0, 3, 5, and 7 post-embedding, endothelial outgrowth surrounding each explant was imaged by brightfield microscopy using a Revolve Microscope (Echo Laboratories, San Diego, CA, USA) and assessed qualitatively.

### Isolation and Culture of Primary Mouse Choroidal Endothelial Cells

After 7 days of incubation, the Matrigel embedding the RPE/choroid/sclera explants from WT and FXR^−/−^ mice was dissolved by Collagenase/Dispase (Sigma-Aldrich Collagenase/Dispase, USA) at 37°C for 2 hours, with repeated pipetting to separate the sprouts. The digested cell suspension from each well was transferred into a 50 mL centrifuge tube. The cells were then filtered through a 70- µm cell strainer and enriched for endothelial cells using CD31-positive magnetic-activated cell sorting (MACS). Endothelial cell identity was verified by the expression of endothelial markers, including CD31 and VE-cadherin. The isolated CD31+ CECs were seeded onto culture plates and maintained in EGM-2 medium supplemented with endothelial growth factors and 10% FBS at 37°C and 5% CO . Cells from early passages (P2– P5) were used for all experiments.

### Endothelial Tube Formation Assay

Primary mouse choroidal endothelial cells (CECs) were isolated from WT and FXR^−/−^ mice as described previously^30^ and exposed to hypoxic conditions (2% O ) for 6 hours in a humidified chamber maintained at 37°C. Hypoxic preconditioning was used to mimic the proangiogenic environment of the CNV lesion and sensitize endothelial angiogenic responses. After 6 hours of hypoxic exposure, cells were collected and used to perform the tube formation assay. Matrigel (354234, Corning, Corning, NY, USA) was thawed overnight at 4°C, and 80 µL was added to each well of a 96-well tissue culture plate. Plates were incubated at 37°C for at least 30 minutes to allow solidification. The CECs collected from the hypoxia chamber were counted and seeded onto the Matrigel-coated wells at 1.2 × 10 cells/well. Cells were incubated at 37°C, and tube formation was assessed beginning at 3 hours and every 0.5–1 hour thereafter.

Images were captured at 6 hours using phase-contrast microscopy at 20× magnification. Cumulative tube length and tube area were analyzed by NIH ImageJ software.

### Fluorescein Angiography

FA was performed to evaluate choroidal neovascularization (CNV) leakage and lesion progression following laser-induced injury. Mice were anesthetized with xylazine hydrochloride (10 mg/kg) and ketamine hydrochloride (100 mg/kg). Pupils were dilated with 1% tropicamide ophthalmic solution (Henry Schein, NY, USA). Then 10% fluorescein sodium (Nexus Pharmaceuticals; NDC 14789-0122-05) was injected intraperitoneally. Fundus fluorescein images were acquired using a Micron IV retinal imaging microscope immediately after fluorescein injection and continuously for approximately 5 minutes. Imaging was performed at 7, 14, and 21 days post-laser injury. CNV lesion morphology was assessed by FA, with images qualitatively evaluated by investigators masked to genotype.

### Hematoxylin and Eosin Staining

Mice were euthanized at 7, 14, and 21 days after laser-induced CNV. Eyes were enucleated, embedded in optimal cutting temperature (O.C.T.) compound, and cryosectioned (8–10 µm). Retinal sections were stained with hematoxylin and eosin using standard protocols. Images were obtained using a Leica brightfield microscope.

### Optical Coherence Tomography

OCT was performed to evaluate retinal morphology and choroidal neovascularization (CNV) lesions in vivo^31^. Mice were anesthetized by injection of ketamine (100 mg/kg) and xylazine (10 mg/kg), and pupils were dilated with a topical drop of 1% tropicamide ophthalmic solution, followed by application of GenTeal Lubricant Eye Gel. Posterior segment structure was assessed at the indicated time points in anesthetized mice using a Bioptigen Spectral Domain Ophthalmic Imaging System (SDOIS). Scans centered on the laser-induced lesions were acquired at 7, 14, and 21 days after laser injury according to the manufacturer’s instructions. Hyperreflective subretinal lesions corresponding to CNV were identified on OCT B-scan images. Lesion area was quantified as previously described ^32^ using ImageJ (National Institutes of Health, Bethesda, MD, USA) by manual delineation of lesion boundaries, performed by investigators masked to genotype. Representative OCT B-scans were generated to illustrate disease progression.

### Choroidal Flat-Mounts/Immunostaining

Mice were euthanized 7, 14, and 21 days after laser-induced choroidal neovascularization (CNV), and eyes were immediately enucleated. Eyes were fixed in 10% neutral-buffered formalin for 2 hours at room temperature. After fixation, the anterior segment and neural retina were carefully removed, and the RPE/choroid/sclera complexes were dissected to obtain posterior eyecups. Tissues were permeabilized in phosphate-buffered saline (PBS) containing 0.1% Triton X-100 for 1 hour at room temperature and then blocked with 5% normal serum in PBS for 1 hour.

For immunofluorescence staining, RPE/choroid/sclera flat-mounts were incubated overnight at 4°C with antibodies against α-smooth muscle actin (α-SMA; sc-32251, 1:100; Santa Cruz Biotechnology, Dallas, TX, USA) and collagen I (NB600-408, 1:100; Novus Biologicals, Centennial, CO, USA), along with biotinylated isolectin B4 (IB4; 10 µg/mL, Invitrogen) followed by Texas Red-conjugated avidin D (Vector Laboratories, catalog # A-2006) to visualize neovascular lesions. After primary antibody incubation, tissues were washed three times in PBS for 15 minutes each and then incubated with the appropriate fluorescent secondary antibodies. After final washes, flat-mounts were mounted on glass slides with the scleral side facing downward using Fluoroshield with DAPI mounting medium (Sigma-Aldrich).

Fluorescence images were acquired using a Leica Stellaris laser-scanning confocal microscope (Leica Microsystems, Wetzlar, Germany) with identical imaging settings across all experimental groups. Z-stack images encompassing the entire CNV lesion were collected and used for quantitative analyses. CNV lesion area, α-SMA-positive fibrotic area, and collagen I-positive staining area were quantified using ImageJ software (National Institutes of Health, Bethesda, MD, USA) by investigators masked to genotype. Lesion boundaries were manually delineated, and the corresponding positive-staining areas were calculated and expressed in square micrometers (µm²).

### Immunofluorescence Staining of Retinal Cross-Sections

Mice were euthanized 14 days after laser-induced CNV, and eyes were enucleated and embedded directly in O.C.T. compound without prior fixation. Cryosections (5–8 µm) were cut through the laser lesions and mounted on slides. Sections were fixed in 4% paraformaldehyde for 15 minutes at room temperature, washed in PBS containing 0.1% Tween-20 (PBST), and blocked with universal blocking buffer (Biogenex Laboratories). Sections were then incubated overnight at 4°C with primary antibodies against VE-cadherin (550548, 1:100; BD Biosciences, San Jose, CA, USA) and α-SMA (sc-32251, 1:100; Santa Cruz Biotechnology, Dallas, TX, USA), together with biotinylated IB4 (10 µg/mL, Invitrogen) followed by Texas Red-conjugated avidin D (Vector Laboratories, catalog # A-2006) to label vasculature. After washing in PBST, sections were incubated with the appropriate fluorescent secondary antibodies and mounted with Fluoroshield mounting medium containing DAPI (Sigma-Aldrich). Images were acquired on a Zeiss confocal microscope using identical acquisition settings across groups and both merged and single-channel images were generated.

### Statistics

Western blot densitometry (fold change relative to control, normalized to β-actin) and endothelial tube formation were compared using a Welch-corrected t-test. Neovascular (IB4- positive) area, lesion area (measured by OCT), and fibrotic areas (α-SMA and collagen I) were analyzed by two-way ANOVA (genotype × time) followed by Šídák-adjusted multiple comparisons. Normality was checked using the Shapiro-Wilk test (P > 0.05 for all groups). For the human transcriptomic data, NR1H4 expression across ocular compartments was compared by the Kruskal-Wallis test followed by Dunn’s multiple-comparison test, and differential expression between AMD and control macular RPE/choroid was determined using DESeq2 with Benjamini-Hochberg correction. Results are presented as mean ± SEM; P < 0.05 was considered statistically significant. Statistical analyses were performed in GraphPad Prism version 8, and transcriptomic analyses were performed using DESeq2 in R.

## Results

### FXR is enriched in the RPE/choroid and reduced in human AMD and experimental CNV

Analysis of the GSE135092 human transcriptomic dataset showed that NR1H4 (FXR) was expressed predominantly in the RPE/choroid compared with the neural retina. Mean NR1H4 expression in macular RPE/choroid was approximately 61-fold higher than in macular retina (2.86 vs 0.047 reads per kilobase of transcript per million mapped reads [RPKM]), a distribution that paralleled canonical RPE identity markers including RPE65, BEST1, LRAT, RLBP1, and TYR (Figure 1A, B). NR1H4 was detectable in all macular RPE/choroid samples examined, supporting expression of FXR within the tissue compartment in which CNV develops.

**Figure 1.**
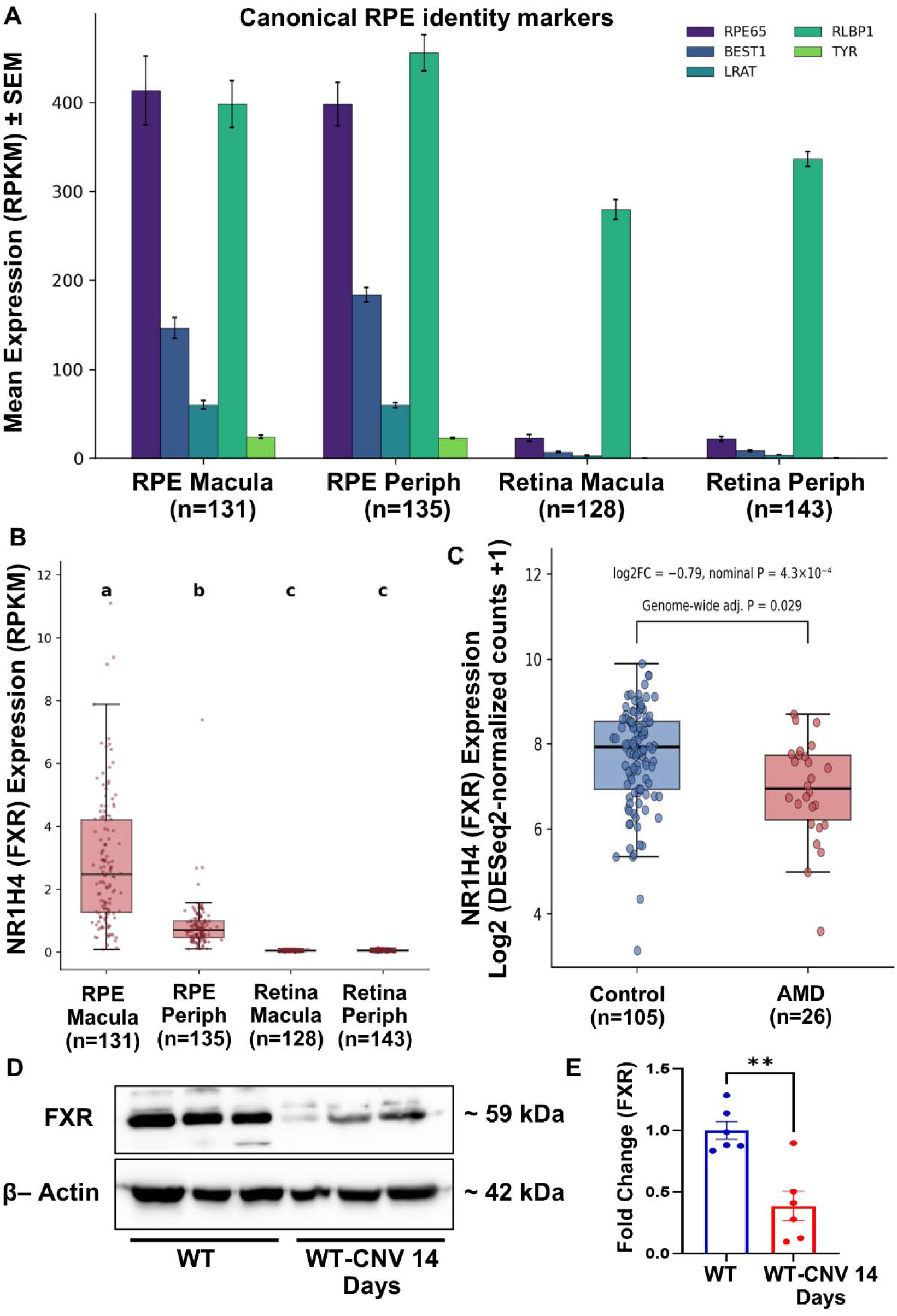
FXR (NR1H4) is expressed in the RPE/choroid and is downregulated in human AMD and mouse laser-induced CNV. (A) Mean expression (RPKM ± SEM) of canonical RPE identity markers (RPE65, BEST1, LRAT, RLBP1, TYR) across four ocular compartments in the human GSE135092 dataset: macular RPE/choroid (n = 131), peripheral RPE/choroid (n = 135), macular retina (n = 128), and peripheral retina (n = 143). Samples from AMD and control donors were pooled to characterize tissue-level expression independent of disease status. RLBP1 is expressed in both the RPE and Müller glia, accounting for its signal in the neural retina. (B) NR1H4 (FXR) expression (RPKM) across the same four compartments, shown as median and interquartile range with individual samples overlaid. Groups not sharing a lowercase letter differ significantly (Kruskal-Wallis test, P < 0.0001; Dunn’s multiple-comparison test, all pairwise comparisons P < 0.0001 except retina macula vs retina periphery, which did not differ). (C) NR1H4 expression in AMD versus control macular RPE/choroid, shown as log2(DESeq2-normalized counts + 1); box, median and interquartile range; points, individual samples. NR1H4 was significantly reduced in AMD (log2 fold-change = −0.79; nominal P = 4.3 × 10 ; genome-wide Benjamini-Hochberg-adjusted P = 0.029; DESeq2; n = 105 control, 26 AMD). (D) Representative Western blot of FXR and β-actin (loading control) in RPE/choroid lysates from untreated wild-type (WT) mice and WT mice 14 days after laser-induced CNV (WT-CNV). (E) FXR protein levels were measured and normalized to β-actin, and results were expressed as fold change compared with WT controls. FXR was significantly reduced in WT-CNV tissue (mean difference −0.61 ± 0.14; Welch- corrected two-tailed t-test, P = 0.0023; n = 6 per group). Data are presented as mean ± SEM. **P < 0.01.

Within macular RPE/choroid, NR1H4 expression was significantly lower in AMD- affected samples than in controls (log2 fold-change = −0.79; nominal P = 4.3 × 10 ; genome- wide Benjamini-Hochberg-adjusted P = 0.029; Figure 1C). To determine whether a similar change occurs during experimental neovascular injury, FXR protein was measured in mouse RPE/choroid 14 days after laser-induced CNV. FXR protein in WT-CNV tissue was reduced to approximately 40% of untreated WT control levels (mean difference −0.61 ± 0.14; Welch- corrected two-tailed t-test, P = 0.0023; Figure 1D, E). Thus, reduced FXR expression was observed in both human AMD RPE/choroid and the mouse CNV model.

### FXR deficiency enhances choroidal angiogenic activity ex vivo and in endothelial cells

To determine whether FXR loss alters choroidal angiogenic behavior, RPE/choroid/sclera explants from WT and FXR^−/−^ mice were evaluated in an ex vivo sprouting assay. Minimal outgrowth was present at day 0 in both genotypes. Endothelial sprouting became evident by day 3 and increased through days 5 and 7. Across the time course, FXR^−/−^ explants showed more extensive radial outgrowth and denser microvascular sprouting than WT explants, with the difference most apparent by day 7 (Figure 2A, B). Because sprouting was assessed qualitatively, these observations were considered supportive evidence of enhanced angiogenic activity in FXR- deficient choroidal tissue.

**Figure 2.**
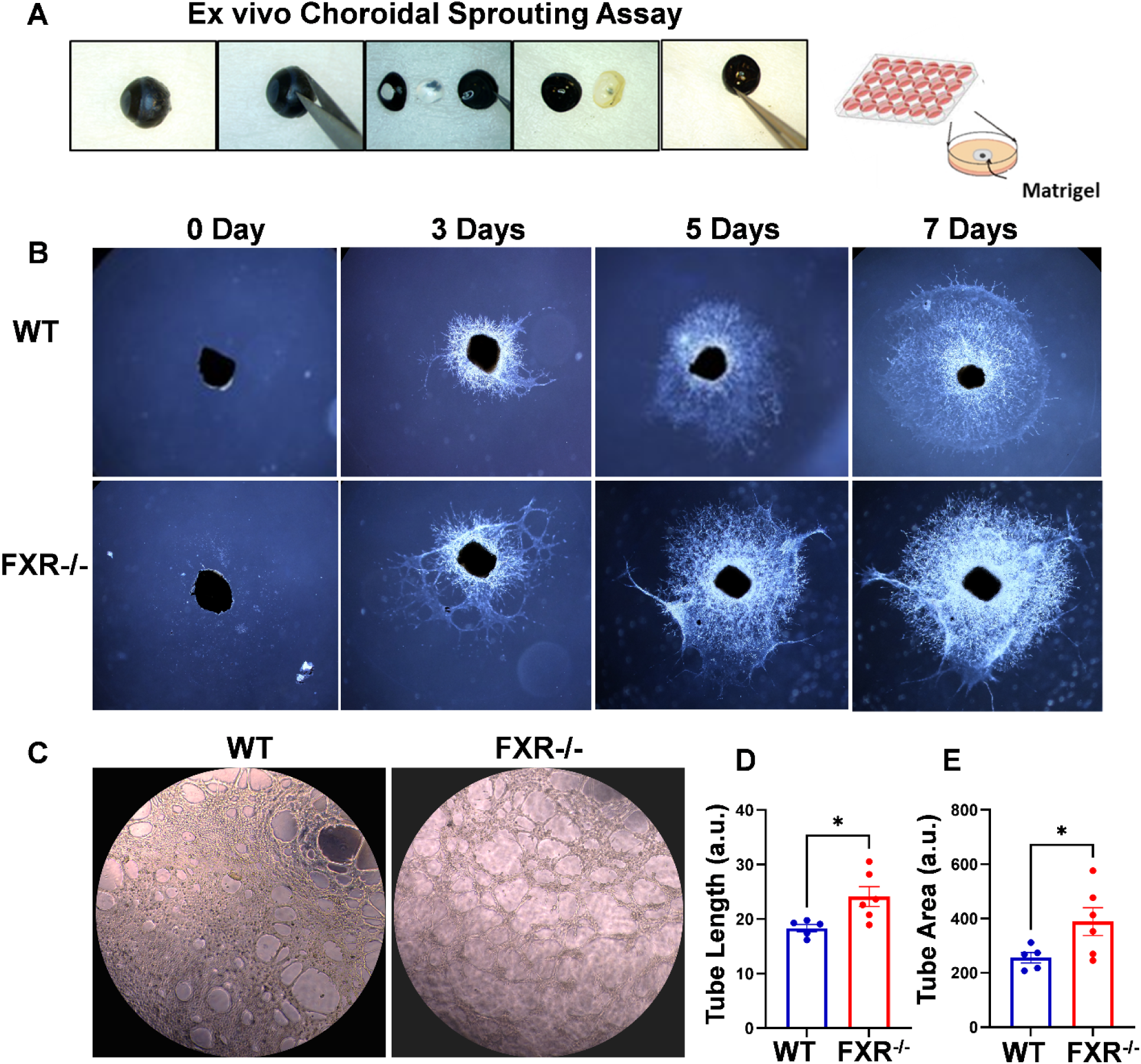
FXR deficiency enhances choroidal endothelial sprouting and tube formation. (A) Schematic of the ex vivo choroidal sprouting assay: RPE/choroid/sclera explants were dissected and embedded in Matrigel. (B) Representative brightfield images of choroidal explants from wild-type (WT) and FXR^−/−^ mice at 0, 3, 5, and 7 days after embedding, showing progressive endothelial sprouting. FXR^−/−^ explants displayed more extensive sprouting outgrowth than WT explants across the time course. (C) Representative images of endothelial tube formation by primary choroidal endothelial cells isolated from WT and FXR^−/−^ mice. (D, E) Quantification of (D) tube length and (E) tube area. FXR^−/−^ cells formed networks with significantly greater tube length (WT 18.4 vs FXR^−/−^ 24.1; P = 0.0228) and tube area (WT 255.8 vs FXR^−/−^ 389.1; P = 0.0492) than WT cells (unpaired t-test with Welch’s correction; n = 5 WT, 6 FXR^−/−^). Data are presented as mean ± SEM. *P < 0.05.

Consistent with the explant findings, primary choroidal endothelial cells isolated from FXR^−/−^ mice formed more extensive capillary-like networks on Matrigel than WT cells. FXR^−/−^ endothelial cells exhibited significantly greater total tube length (WT 18.4 vs FXR^−/−^ 24.1 arbitrary units [a.u.]; P = 0.0228) and tube area (WT 255.8 vs FXR^−/−^ 389.1 a.u.; P = 0.0492) (Figure 2C–E). These findings indicate that loss of FXR enhances endothelial network formation under the conditions tested.

### FXR deficiency increases CNV lesion severity, vascular leakage, and outer retinal disruption

The experimental timeline for laser-induced CNV and subsequent imaging is summarized in Figure 3A. The effect of FXR deficiency on CNV progression was next assessed by serial fundus imaging and FA at 7, 14, and 21 days after laser injury. In WT mice, focal lesions were evident by day 7, were most prominent at day 14, and appeared reduced by day 21. FXR^−/−^ eyes showed larger and more prominent lesions throughout the same interval (Figure 3B).

**Figure 3.**
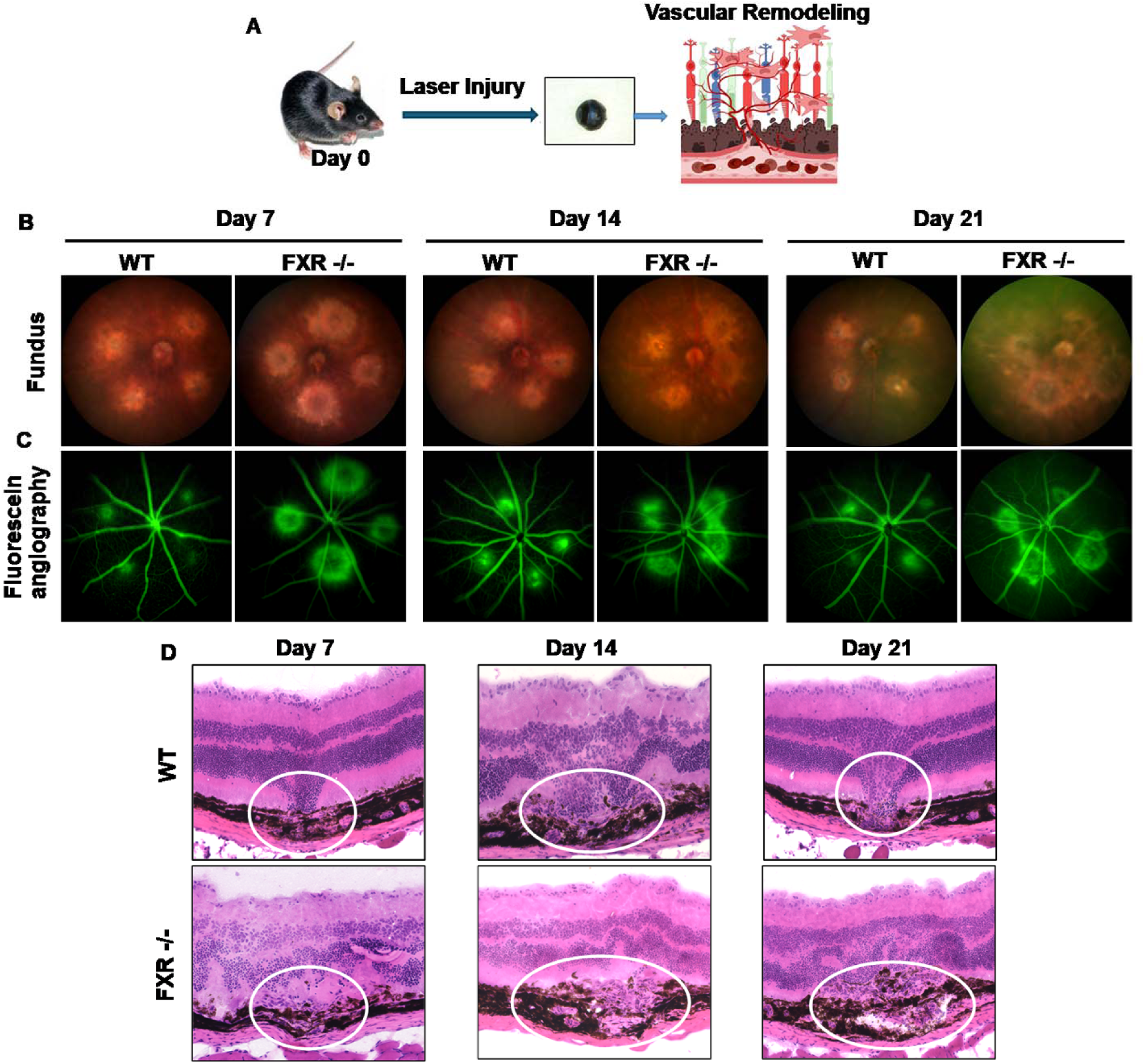
FXR deficiency exacerbates CNV lesion formation, vascular leakage, and retinal structural damage. (A) Schematic of the experimental design: laser photocoagulation was applied to induce choroidal neovascularization (CNV), and lesions were evaluated during the subsequent vascular remodeling phase. (B) Representative fundus photographs of wild-type (WT) and FXR^−/−^ eyes at 7, 14, and 21 days after laser injury. (C) Representative fluorescein angiography (FA) images of the same eyes, showing hyperfluorescence at sites of vascular leakage; FXR^−/−^ eyes displayed greater hyperfluorescence than WT across the time course. (D) Representative hematoxylin and eosin (H&E)-stained retinal cross-sections of WT and FXR^−/−^ eyes at 7, 14, and 21 days after laser injury. Lesion sites are outlined; FXR^−/−^ retinas showed larger and more disorganized subretinal lesions with greater disruption of the RPE and outer retina than WT. Images are representative of 5 animals per group.

FA demonstrated a similar temporal pattern. WT eyes showed localized hyperfluorescence at laser lesions that increased through day 14 and decreased by day 21. In contrast, FXR^−/−^ eyes exhibited larger and more intense hyperfluorescent lesions at each time point, with persistent leakage at day 21 (Figure 3C). These qualitative imaging findings were accompanied by greater structural disruption in FXR-deficient eyes.

H&E-stained retinal sections confirmed localized subretinal lesions at the laser sites in both genotypes. WT lesions were most prominent at day 14 and appeared more contained by day 21, whereas FXR^−/−^ eyes showed larger, less organized lesions with greater disruption of the RPE and outer retinal architecture across the time course (Figure 3D).

### Quantitative imaging confirms greater and more persistent CNV lesion burden in FXR^−/−^ mice

Quantification of IB4-positive neovascular complexes in RPE/choroid/sclera flat-mounts showed that lesion area increased from day 7 to day 14 in both genotypes and then declined by day 21 (Figure 4A, B). Neovascular area rose significantly from day 7 to day 14 in both WT (P = 0.011) and FXR^−/−^ (P = 0.016) mice and declined significantly from day 14 to day 21 in both genotypes (WT, P = 0.015; FXR / , P < 0.0001). FXR^−/−^ mice had significantly larger IB4- positive lesions than WT mice at day 7 (P < 0.0001), day 14 (P < 0.0001), and day 21 (P = 0.0078). Despite declining from its day 14 peak, FXR^−/−^ neovascular area remained significantly greater than WT at day 21, indicating persistence of an increased neovascular burden in the absence of FXR.

**Figure 4.**
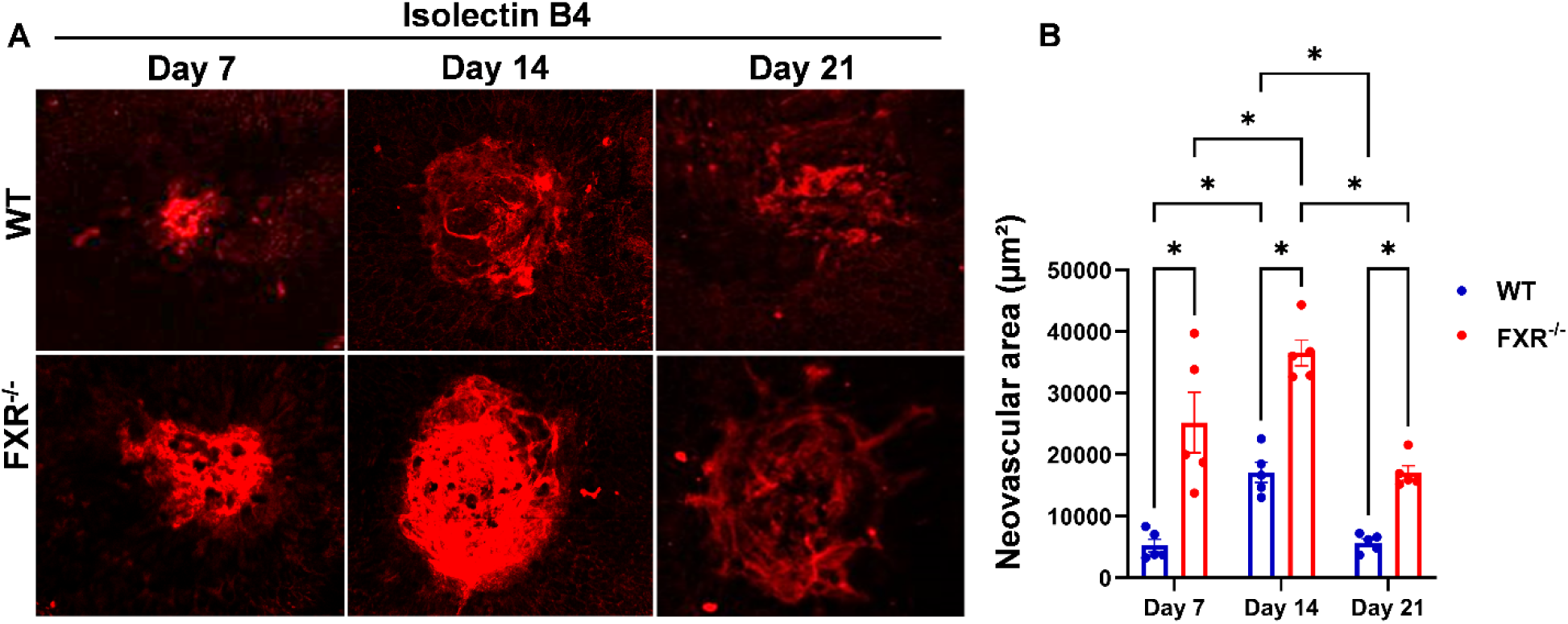
FXR deficiency increases neovascular area in RPE/choroid flat-mounts. (A) Representative IB4 (IB4)-stained RPE/choroid/sclera flat-mounts showing CNV lesions in wild-type (WT) and FXR^−/−^ mice at 7, 14, and 21 days after laser injury. (B) Quantification of IB4-positive neovascular area (µm²). Data were analyzed by two-way ANOVA (genotype × time): genotype, F(1, 24) = 75.24, P < 0.0001; time, F(2, 24) = 22.59, P < 0.0001; interaction, F(2, 24) = 2.05, P = 0.15; followed by Šídák’s multiple-comparison test. FXR^−/−^ mice had greater neovascular area than WT at day 7 (P < 0.0001), day 14 (P < 0.0001), and day 21 (P = 0.0078). Within both genotypes, neovascular area increased from day 7 to day 14 (WT, P = 0.011; FXR^−/−^, P = 0.016) and declined from day 14 to day 21 (WT, P = 0.015; FXR^−/−^, P < 0.0001). Data are presented as mean ± SEM; each point represents the average of 2–4 lesions from one animal (n = 5 animals per genotype per time point). *P < 0.05; exact p-values are given above.

OCT provided an independent in vivo measure of lesion burden. Hyperreflective subretinal lesions were identified at the laser sites at days 7, 14, and 21 (Figure 5A). Quantified OCT lesion area was significantly greater in FXR^−/−^ than WT mice at all three time points: day 7 (P < 0.0001), day 14 (P < 0.0001), and day 21 (P < 0.0001) (Figure 5B). Representative B-scans also showed greater disruption of the outer retinal architecture surrounding FXR^−/−^ lesions. The genotype difference was consistent across time, with no significant genotype × time interaction (P = 0.09), indicating that FXR^−/−^ lesions were larger than WT at every time point examined. Together with the flat-mount data, these OCT measurements confirm that FXR deficiency increases CNV lesion size.

**Figure 5.**
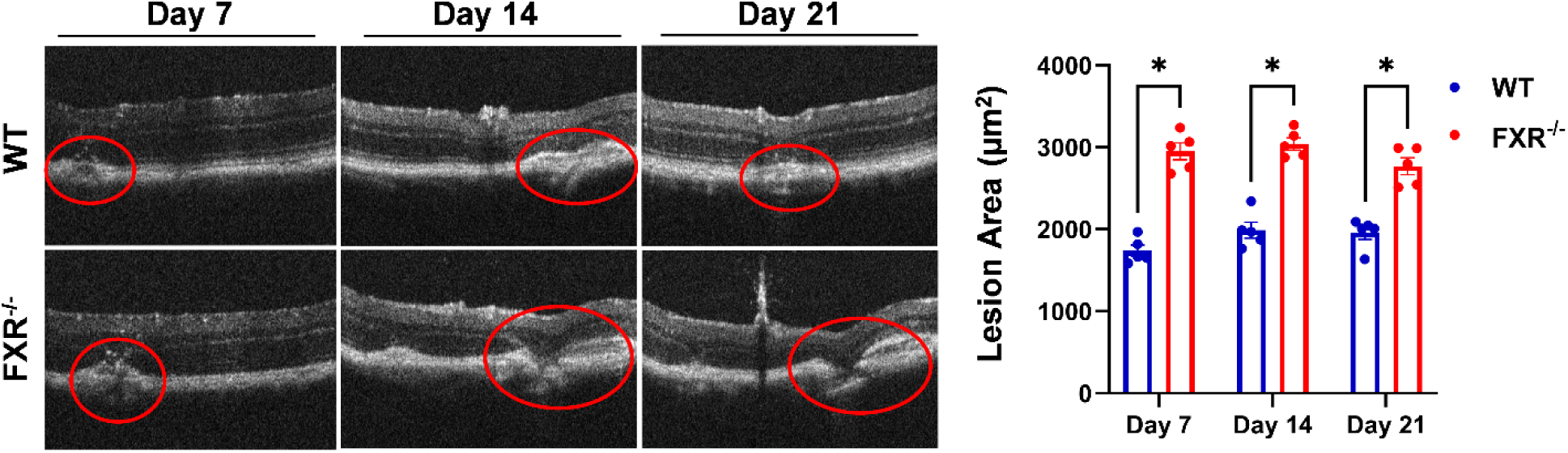
FXR deficiency increases retinal lesion area following laser-induced CNV. (A) Representative spectral-domain OCT B-scans of wild-type (WT) and FXR^−/−^ retinas at 7, 14, and 21 days after laser injury, showing hyperreflective subretinal lesions at the laser sites. (B) Quantification of OCT-measured lesion area (µm²). Data were analyzed by two-way ANOVA (genotype × time): genotype, F(1, 24) = 198.2, P < 0.0001; time, F(2, 24) = 2.19, P = 0.13; interaction, F(2, 24) = 2.61, P = 0.09; followed by Šídák’s multiple-comparison test. FXR^−/−^ mice had greater lesion area than WT mice at each time point (day 7, day 14, and day 21; all P < 0.0001). Data are presented as mean ± SEM; each point represents the average of 4–6 lesions from one animal (n = 5 animals per genotype per time point). *P < 0.05; exact p-values are given above.

### FXR deficiency promotes persistent fibrotic and mesenchymal remodeling after CNV injury

To determine whether FXR also influences the fibrovascular remodeling phase of CNV, RPE/choroid/sclera flat-mounts were analyzed for collagen I and α-smooth muscle actin (α- SMA) at days 7, 14, and 21 after laser injury (Figure 6). Collagen I-positive area was significantly greater in FXR^−/−^ than WT lesions at all three time points: day 7 (P = 0.046), day 14 (P < 0.0001), and day 21 (P = 0.0024) (Figure 6A, B). Collagen I did not change significantly over time in WT mice, whereas in FXR^−/−^ mice it rose from day 7 to a peak at day 14 (P < 0.0001) and then declined by day 21 (P < 0.0001) while remaining significantly greater than WT. The significant genotype × time interaction (P < 0.0001) reflected the markedly greater collagen I accumulation in FXR^−/−^ lesions at the day 14 peak, indicating sustained extracellular matrix deposition in the absence of FXR during the remodeling phase.

**Figure 6.**
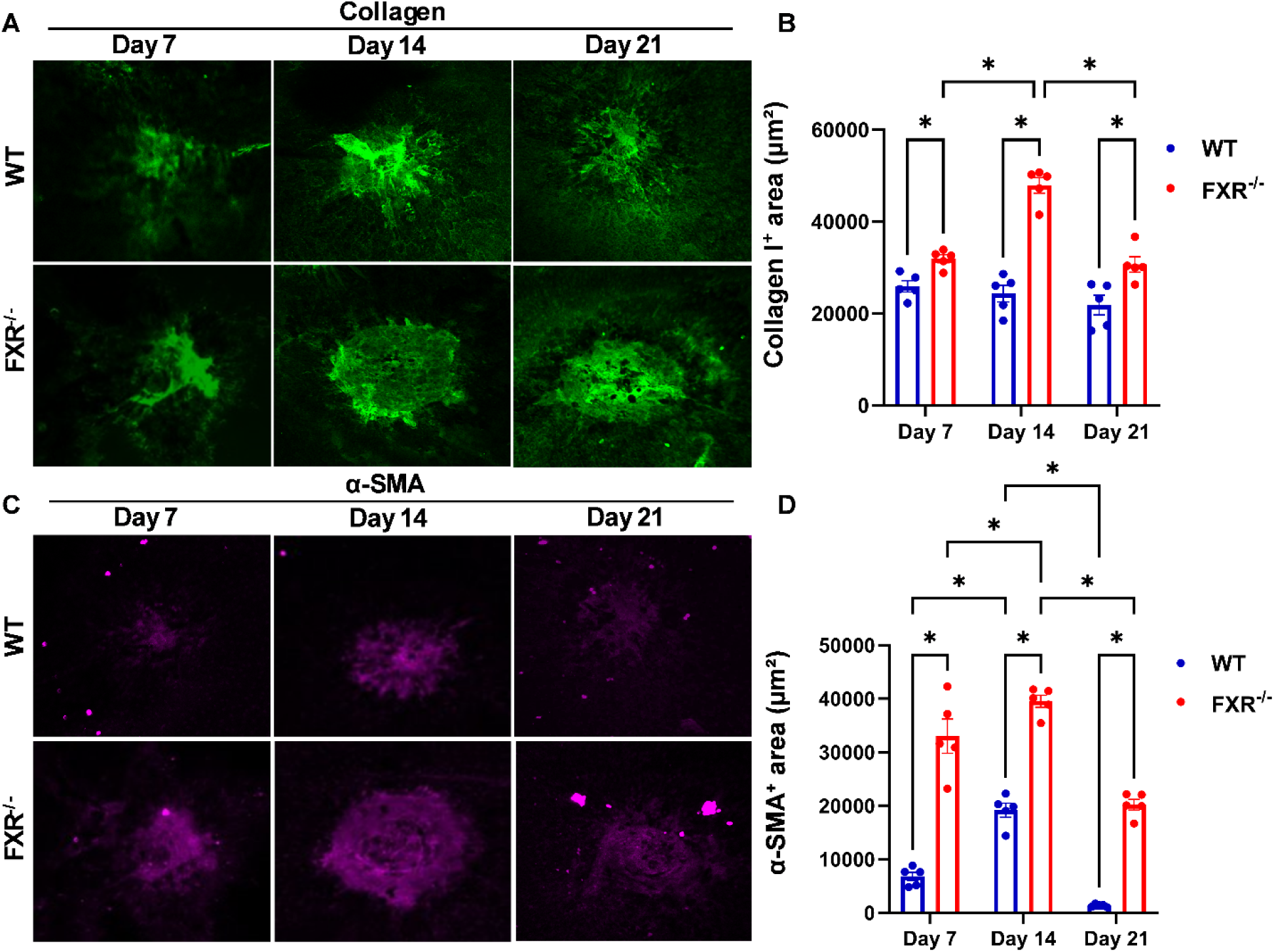
FXR deficiency enhances fibrotic remodeling during CNV. (A) Representative collagen I-stained RPE/choroid/sclera flat-mounts from wild-type (WT) and FXR^−/−^ mice at 7, 14, and 21 days after laser injury. (B) Quantification of collagen I-positive area (µm²). Data were analyzed by two-way ANOVA (genotype × time): genotype, F(1, 24) = 92.94, P < 0.0001; time, F(2, 24) = 19.63, P < 0.0001; interaction, F(2, 24) = 16.84, P < 0.0001; followed by Šídák’s multiple-comparison test. Collagen I positive area was significantly greater in FXR^−/−^ than WT mice at day 7 (P = 0.046), day 14 (P < 0.0001), and day 21 (P = 0.0024). In FXR^−/−^ mice, collagen I increased from day 7 to day 14 (P < 0.0001) and declined from day 14 to day 21 (P < 0.0001), whereas it did not change significantly over time in WT mice. (C) Representative α-smooth muscle actin (α-SMA)-stained flat-mounts from WT and FXR^−/−^ mice at the same time points. (D) Quantification of α-SMA positive (fibrotic) area (µm²). Data were analyzed by two-way ANOVA (genotype × time): genotype, F(1, 24) = 286.7, P < 0.0001; time, F(2, 24) = 69.33, P < 0.0001; interaction, F(2, 24) = 3.07, P = 0.065; followed by Šídák’s multiple-comparison test. α-SMA positive area was significantly greater in FXR^−/−^ than WT mice at day 7, day 14, and day 21 (all P < 0.0001). α-SMA positive area declined significantly from day 14 to day 21 in both genotypes (both P < 0.0001), but FXR^−/−^ α-SMA positive area remained significantly greater than WT at day 21. Data are presented as mean ± SEM; each point represents the average of 2–4 lesions from one animal (n = 5 animals per genotype per time point). *P < 0.05; exact p-values are given above.

A similar pattern was observed for α-SMA. α-SMA-positive area was significantly greater in FXR^−/−^ than in WT mice at every time point, day 7, day 14, and day 21 (all P < 0.0001) (Figure 6C, D). α-SMA-positive area peaked at day 14 and declined by day 21 in both genotypes (day 14 vs day 21: WT, P < 0.0001; FXR^−/−^, P < 0.0001); however, whereas WT α- SMA fell to near-baseline levels, FXR^−/−^ α-SMA remained markedly and significantly elevated above WT at day 21 (P < 0.0001), indicating prolonged myofibroblast and mesenchymal activation in the absence of FXR.

### FXR deficiency is associated with increased TGF-β1 and mesenchymal activation in CNV lesions

Because the sustained α-SMA response suggested persistent mesenchymal activation, lesions were further examined at day 14 for endothelial/mesenchymal marker distribution and TGF-β1 expression. Immunofluorescence showed increased α-SMA labeling and more disorganized vascular architecture in FXR^−/−^ CNV lesions compared with WT-CNV lesions (Figure 7A, B). Western blot analysis demonstrated a 2.4-fold increase in TGF-β1 protein in FXR^−/−^ RPE/choroid relative to WT-CNV tissue (P = 0.0286; n = 6 per group; Figure 7C, D). These data associate FXR deficiency with enhanced TGF-β1 signaling and mesenchymal activation during the peak remodeling phase, without establishing endothelial lineage conversion.

**Figure 7.**
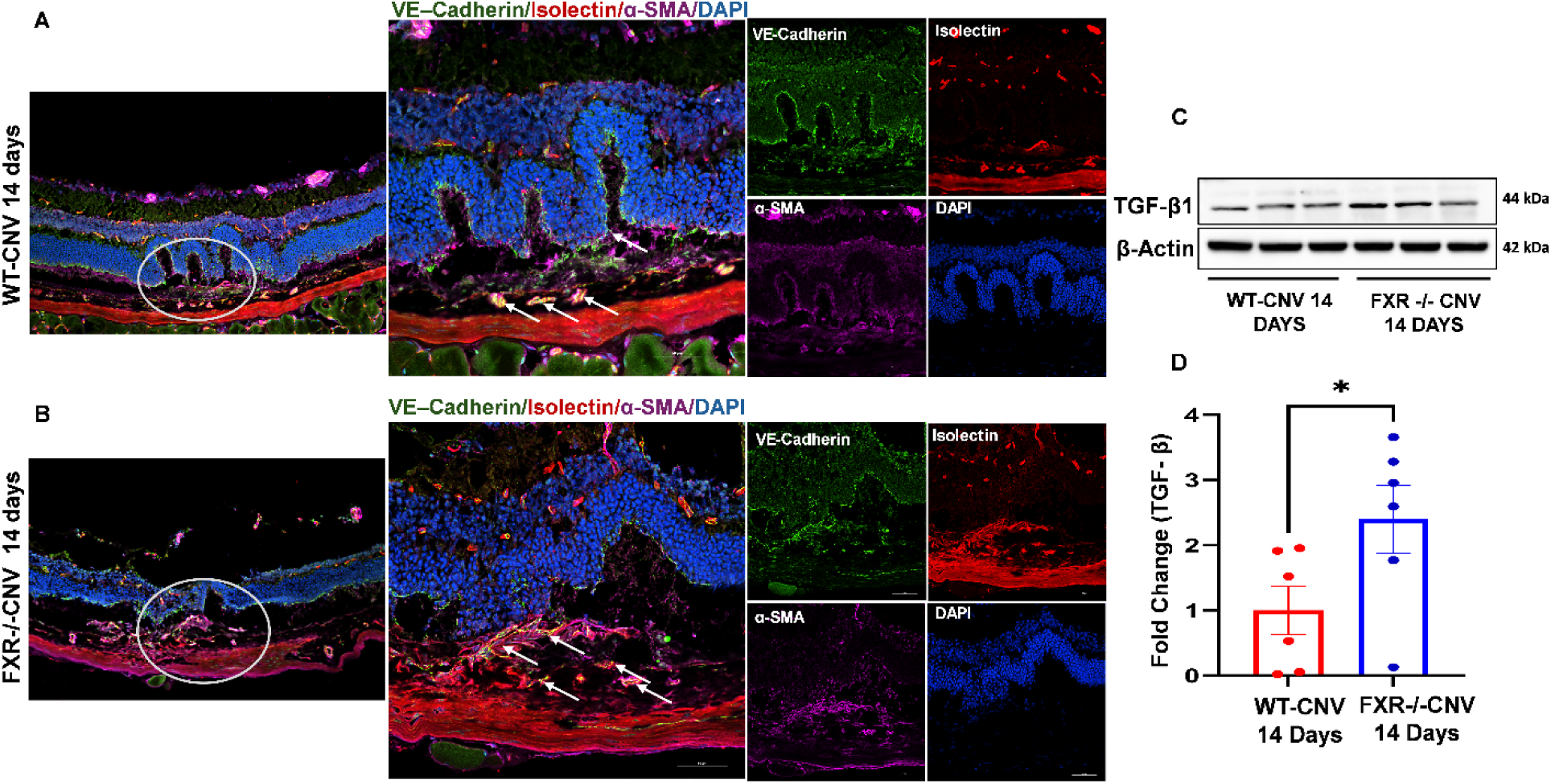
FXR deficiency increases TGF-β1 expression and mesenchymal activation in CNV lesions. (A, B) Representative immunofluorescence of RPE/choroid cross-sections from (A) WT-CNV and (B) FXR^−/−^ CNV mice at 14 days after laser injury, co-stained for VE-cadherin (endothelial marker, green), IB4 (vasculature, red), α-SMA (myofibroblasts, magenta), and DAPI (nuclei, blue); merged and single-channel images are shown, and the lesion area is outlined. FXR^−/−^ lesions showed increased α-SMA labeling and disorganized vasculature compared with WT (arrows). (C) Western blot and (D) quantification of TGF-β1 (normalized to β-actin) in RPE/choroid lysates from WT-CNV and FXR^−/−^ CNV mice at 14 days. TGF-β1 was increased in FXR^−/−^ relative to WT-CNV (2.4-fold; unpaired t-test with Welch’s correction, P = 0.0286; n = 6 per group). Data are presented as mean ± SEM. *P < 0.05.

## Discussion

The present study identifies FXR as an endogenous regulator of choroidal neovascularization and lesion remodeling. Human transcriptomic analysis demonstrated enrichment of NR1H4 in the RPE/choroid and a significant reduction in AMD-affected macular RPE/choroid, while FXR protein was similarly reduced in mouse RPE/choroid at day 14 after laser-induced CNV. Functional loss of FXR was associated with greater choroidal sprouting, enhanced endothelial network formation, increased CNV lesion size and vascular leakage, greater outer retinal disruption, and persistent fibrotic remodeling. Together, these findings support a protective role for FXR in restraining both the angiogenic and fibrotic components of CNV.

Our analysis of a publicly available human transcriptomic dataset (GSE135092) ^27^ showed that NR1H4 is expressed in and enriched in the RPE/choroid complex, the tissue in which CNV originates, relative to neural retina. Importantly, NR1H4 expression was significantly reduced in AMD-affected macular RPE/choroid as compared to controls. Notably, to our knowledge, no genome-wide AMD association has been reported at the NR1H4 locus, indicating that any contribution of FXR to AMD likely acts at the level of expression or function rather than inherited genetic risk.

In the mouse model, FXR expression was reduced at day 14, when neovascular and fibrotic activity were most prominent. Similar reductions in FXR have been described in other fibrotic tissues, where diminished FXR signaling is associated with enhanced inflammatory and profibrotic responses ^33^. The present data do not establish why FXR expression falls during CNV, but they suggest that loss of this regulatory pathway may accompany a microenvironment that favors continued vascular and mesenchymal activation.

The endothelial findings provide one potential explanation for the larger CNV phenotype in FXR-deficient mice. FXR^−/−^ choroidal explants showed greater vascular outgrowth, and primary FXR^−/−^ choroidal endothelial cells formed networks with increased tube length and area. Although the explant assay was qualitative, the concordance between ex vivo sprouting and quantitative endothelial tube formation supports a role for FXR in limiting angiogenic behavior within the choroidal vascular compartment. The vascular actions of FXR are likely context- dependent, however, because FXR activation has been reported to exert different effects on endothelial migration and network formation in other vascular beds ^34^. Defining the downstream endothelial targets of FXR in the choroid will therefore be important.

The longitudinal in vivo data further indicate that FXR influences more than the initial angiogenic response. Fundus imaging, FA, flat-mount analysis, and OCT consistently showed a greater lesion burden in FXR^−/−^ mice from day 7 through day 21. In WT mice, lesion size and leakage increased toward day 14 and then partially regressed by day 21, whereas FXR^−/−^ lesions remained larger during this remodeling phase. The persistence of increased lesion area at day 21 suggests that FXR contributes to containment and/or resolution of CNV after the initial vascular growth phase, rather than acting solely at lesion initiation.

The fibrotic data show that FXR deficiency enhances both collagen deposition and myofibroblast activation during CNV. Collagen I-positive area was significantly greater in FXR^−/−^ than WT lesions at all three time points (day 7, 14, and 21); in FXR^−/−^ mice it peaked at day 14 and declined by day 21 but remained above WT, whereas WT collagen I changed little over time. α-SMA-positive area was likewise greater in FXR^−/−^ than WT at every time point; although α-SMA peaked at day 14 and declined by day 21 in both genotypes, FXR^−/−^ α-SMA remained markedly elevated at day 21, when the WT response had largely resolved. Together, these data indicate a sustained fibrotic and myofibroblast response in the absence of FXR. The accompanying increase in TGF-β1 at day 14 provides a plausible link to this phenotype, because TGF-β signaling is a major driver of myofibroblast differentiation and has been implicated in epithelial-to-mesenchymal transition/endothelial-to-mesenchymal transition (EMT/EndoMT)- associated subretinal fibrosis ^11, 35^. However, the current marker-based data do not demonstrate endothelial lineage conversion, and lineage-tracing studies will be required to determine whether EndoMT directly contributes to the FXR-deficient phenotype.

Collectively, the results support a model in which FXR helps maintain RPE/choroidal homeostasis during injury by constraining endothelial activation during lesion growth and limiting sustained mesenchymal activation during remodeling. This study has several limitations. First, the global FXR knockout model cannot differentiate the roles of endothelial, RPE, immune, and other cells; cell-specific approaches are needed. Second, increased TGF-β1 and persistent α-SMA suggest mesenchymal activation, but causality between FXR loss and the TGF-β/SMAD pathway is not confirmed, and EndoMT is not established. Third, only male mice were studied; whether FXR exerts sex-dependent effects on CNV progression remains untested. Fourth, the human transcriptomic analysis from bulk RPE/choroid cannot identify the cell type responsible for reduced NR1H4 or prove causation. In addition, the AMD samples in this dataset comprise pooled AMD stages, including both atrophic and neovascular forms, so the reduction in NR1H4 reflects an association with AMD broadly rather than with choroidal neovascularization specifically. Our ongoing studies will subsequently determine which of these pathways are directly controlled by FXR in endothelial cells, RPE, and immune cells within the CNV lesion, and whether pharmacologic FXR activation can reduce established neovascular and fibrotic disease. Such studies will be particularly important before positioning FXR-directed therapy as an adjunct to VEGF-targeted treatment.

## Author Contributions

CH: Methodology, Investigation, Formal Analysis, Data Curation, Writing – Review and Editing. MAN: Methodology, Investigation, Data Curation, Writing – Review and Editing. AS: Data Curation, Validation, Writing – Review and Editing. AO: Data Curation, Methodology, Writing – Review and Editing. BB: Data Curation, Methodology, Writing – Review and Editing.GS: Data Curation, Methodology, Writing – Review and Editing. PMM: Supervision, Resources, Funding, Project Administration, Writing – Review and Editing. RNJ: Supervision, Resources, Funding, Project Administration, Writing – Review and Editing. MCT: Conceptualization, Investigation, Formal Analysis, Supervision, Project Administration, Funding, Writing- Original Draft, Writing – Review and Editing. All authors contributed to the article and approved the submitted version.

## Ethics Statement

The animal study was approved by the Institutional Animal Care and Use Committee(s) of Augusta University and Meharry Medical College. All procedures were conducted in accordance with institutional requirements and the ARVO Statement for the Use of Animals in Ophthalmic and Vision Research. Publicly available human gene expression data were obtained from the GEO database; all datasets were previously published and de-identified, and therefore no additional institutional review board approval was required for this study.

## Data Availability Statement

The data supporting the findings of this study are included in the article and/or its supplementary material. Additional information is available from the corresponding author upon reasonable request.

## Conflict of Interest Statement

The authors declare no competing interests related to this work.

## Generative AI Disclosure

Generative AI (Grammarly and Anthropic Claude) was used for language editing and manuscript preparation. The authors reviewed and revised all AI-assisted text and take full responsibility for the scientific content, interpretation, accuracy, and final manuscript.

## Funding

The authors’ laboratory is funded by the EY034568 grant and the Chan Zuckerberg Initiative (CZIF2022-007043) accelerate precision health program to MCT, EY035336 to RNJ, and EY033264 to RNJ & PMM. Student trainees were supported by GM144927, HL007737, AI007281, and UC2GM162929. CH is a recipient of the William Townsend Porter Physiology Development Fellowship.

## Acknowledgments

The authors thank Dr. Olga Korolkova, Research Staff Scientist at the Meharry Medical College Core Facilities, for her valuable technical expertise and assistance in acquiring retinal flat-mount images used in this study. The Meharry Medical College Core Facilities are supported by NIH Grants MD007586, CA163069, and S10RR025497

## Notes

### Competing Interest Statement

The authors have declared no competing interest.

